# Bcl-x_L_ inhibition enhances Dinaciclib-induced cell death in soft-tissue sarcomas

**DOI:** 10.1101/400986

**Authors:** Santi Rello-Varona, Miriam Fuentes-Guirado, Roser López-Alemany, Aida Contreras-Pérez, Núria Mulet-Margalef, Silvia García-Monclús, Oscar M. Tirado, Xavier García del Muro

**Author notes:** Corresponding authors: Santi Rello-Varona. Laboratori d’Oncologia Molecular, Institut d’Investigació Biomèdica de Bellvitge (IDIBELL). Hospital Duran i Reynals - 3^a^ Planta, Avda Gran Via 199. 08908 L’Hospitalet de Llobregat, Barcelona, Spain. Òscar M. Tirado. Laboratori d’Oncologia Molecular, Institut d’Investigació Biomèdica de Bellvitge (IDIBELL). Hospital Duran i Reynals - 3^a^ Planta, Avda Gran Via 199. 08908 L’Hospitaletde Llobregat, Barcelona, Spain. Xavier Garcia del Muro. Servei d’Oncologia Mèdica-Divisió D, Institut Català d’Oncologia L’Hospitalet. Hospital Duran i Reynals, Avda Gran Via 199. 08908 L’Hospitalet de Llobregat, Barcelona, Spain. **Conflict of Interests:** Authors declare no conflict of interests. **Financial support:** We thank CERCA Programme / *Generalitat de Catalunya* for institutional support. S. Rello-Varona was a Marie Curie COFUND-Beatriu De PinÒs Researcher (The European Union 7^th^ Framework Program for R+D and *Generalitat de Catalunya*, Department for Economy and Knowledge). S. Rello-Varona, M. Fuentes-Guirado and S. García-Monclús were funded by *Fundación Alba Pérez lucha contra el cáncer infantil*, through a grant endorsed to O.M. Tirado. This work was funded by *Instituto de Salud Carlos III: Acción Estratégica de Salud* (grants PI12-01908 and PI15-00234 endorsed to X. García del Muro) and co-funded by European Regional Development Fund (ERDF), a way to build Europe.

## Abstract

Soft-tissue sarcomas (STS) are an uncommon and heterogeneous group of malignancies that result in high mortality. Metastatic STS have very bad prognosis due to the lack of effective treatments. Dinaciclib is a model drug for the family of CDK inhibitors. Its main targets are cell cycle regulator CDK1 and protein synthesis controller CDK9. We present data supporting Dinaciclib ability to inactivate *in vitro* different STS models at nanomolar concentrations. Moreover, the different rhythms of cell death induction allow us to further study into the mechanism of action of the drug. Cell death was found to respond to the mitochondrial pathway of apoptosis. Anti-apoptotic Bcl-x_L_ was identified as the key regulator of this process. Bcl-x_L_ showed a slower decay curve after protein synthesis disruption that in tolerant cell lines was enough to delay apoptosis, as its action cannot be countered by the relative low levels of pro-apoptotic BH3 proteins BIM and PUMA. Combination of Dinaciclib with BH3-mimetics led to quick and massive apoptosis induction *in vitro*, but *in vivo* assessment was prevented due to liver toxicity. Additionally, Bcl-x_L_ inhibitor A-1331852 also synergized with conventional chemotherapy drugs as Gemcitabine. Thus, Bcl-x_L_ targeted therapy arises as a major opportunity to the treatment of STS.

## Introduction

Soft-tissue sarcomas (STS) are a group of tumors derived from mesenchymal precursors with scarce incidence and rich variability (1). Tumors arising from non-epithelial extra-skeletal tissue are generally accounted as STS (2). There has been much improvement in the understanding of the drivers of STS entities: (i) STSs driven by specific chromosome fusions leading to generation of anomalous transcription factors (like *FUS-CHOP* in myxoid liposarcoma) or chromatin remodelers (*SS18-SSX* in synovial sarcoma); (ii) STS that rely on specific mutations (*KIT* in gastrointestinal stromal tumors) and (iii) other STS driven by more complex genomic rearrangements (like leiomyosarcomas or some fibrosarcomas) (3,4).

STS incidence is difficult to estimate due to their variability, and some reports claim that the usual figures could be underestimations (1,5). Clinical prognosis and therapeutic outcome is also highly variable in STS (2). When it is possible, the complete clinical resection make full recovery achievable. However, almost half of the patients will develop metastatic disease. Five-year survival rates are still below 50 %. So, the weight of STS in total cancer death toll is clearly disproportionate to its incidence (4,6).

Thus, STS can benefit for new therapeutic approaches (6). Among the molecular targeted drugs in development, the group of Cyclin-Dependent Kinases (CDKs) inhibitors is one of those concealing major interest (7). CDKs constitute a wide family of Ser/Thr protein kinases that require binding with cyclins to act. This coupling enables a complex panorama of interactions that keep track on the activation/suppression of specific pathways during cell cycle (8). Several CDK inhibitors have been identified and tested *in vitro* as anti-cancer agents (7,9). The initial aim of CDK inhibitor strategy was the disruption of cell cycle sequence-of-events in order to induce cell death (9,10). But it was soon understood that CDKs exert more powerful effects over other processes of cell physiology like transcription regulation, RNA splicing or protein folding (9).

Dinaciclib is a promising CDK inhibitor, extensively proved pre-clinically (11). Its known affinities encompass CDK1 (IC_50_=3 nM), CDK2 (IC_50_=1 nM), CDK5 (IC_50_=1 nM) and CDK9 (IC_50_=4 nM) (12). CDK2 is a key cell cycle regulator in meiosis but its significance in somatic cells is relative (8). CDK5 role is less clear. It has been associated with neuronal metabolism via multiple interactions, including proteins from the cytoskeleton (13). Most studies concerning Dinaciclib activity have been focused on the CDK1 control of mitotic entry and CDK9 regulation of gene transcription (14–16). CDK9-dependent down regulation of anti-apoptotic Bcl-2 family member Mcl-1 is commonly regarded as the main mechanism of action of this drug (17,18). Some Phase I clinical trials (mostly in pediatric leukemia) have also been performed with Dinaciclib. Anti-cancer activity was found to be encouraging, but not sufficient for planning monotherapy treatments. Further use of Dinaciclib is thought to rely on combination therapies (14,19,20).

Combination therapies constitute a hot spot in oncology research. It has become clear its benefits avoiding tumor evolution in favor of drug resistant phenotypes (21). Moreover, combination therapies work better than monotherapy even in the absence of synergistic behavior (22). BH3-mimetics are a new class of anti-cancer drugs particularly interesting for these combinations. They are aimed to disturb the balance of the different proteins of the Bcl-2 family, thus favoring apoptosis triggering (23,24). Alone, BH3-mimetics have been successfully used in chronic lymphocytic leukemia since the FDA approval of Venetoclax (25). BH3-mimetics work better when the cells are already undergoing an apoptotic signaling process that has been compensated by expression or activity changes in the Bcl-2 family of proteins. Cells became addicted to these compensative mechanisms creating then an Achilles’ heel for cancer cells (24). Recent screenings are showing that BH3-mimetics boost the cytotoxic potential of a panoply of chemicals, including CDK9 inhibitors (26).

Our aim in the present study is to seek the suitability of Dinaciclib in a series of STS *in vitro* models as cell death inductor, fully characterizing the cellular response to treatment. We have found that Dinaciclib is capable of inducing cell death as single agent. The cellular context, particularly the Bcl-2 family balance, at every model is decisive for the precise behavior after Dinaciclib incubation. Our data support that Bcl-x_L_ inhibition status is central for treatment tolerance. Moreover, Bcl-x_L_ specific inhibitors synergize with Dinaciclib to avoid such tolerance.

## Materials & Methods

### Cell culture

Cell lines (**Table 1**) were cultured in RPMI-1640 supplemented with 10 % (v/v) heat-inactivated FBS, 1 % (v/v) Penicillin-Streptomycin and 1 % (v/v) HEPES (Gibco). All cell lines were incubated in an AutoFlow NU-4750 Incubator (NuAire) at 37 °C in a humidified atmosphere of 5 % CO_2_ in air. Otherwise specified, plasticware were purchased from Jet BioFil.

**Table 1:** List of cell lines employed in the study.

| Cell line | Origin | Provider |
| --- | --- | --- |
| **SW982** | Synovial Sarcoma | ATCC |
| **1273-99** | Synovial Sarcoma | Dr. Carmen de Torres, *Hospital Sant Joan de Deu* (Esplugues de Llobregat, Spain) |
| **SK-LMS-1** | Leiomyosarcoma | ATCC |
| **SK-UT-1** | Leiomyosarcoma | CLS |
| **402-91** | Myxoid Liposarcoma | Prof. Roberto Mantovani (*Università degli Studi di Milano*, Milan, Italy) |
| **1765-92** | Myxoid Liposarcoma | Prof. Roberto Mantovani |
| **SW872** | Dedifferentiated Liposarcoma | ATCC |
| **HT-1080** | Fibrosarcoma | ATCC |

Authenticity of the cell lines was routinely confirmed by STR profiling analysis done at qGenomics SL (Esplugues de Llobregat, Barcelona, Spain).

Exponential growing cells were used for all experiments. Depending on cell size and proliferation rates, 80,000-180,000 cells were seeded in 12-well plates (or 1-2 million cells in P100 dishes) and allowed to grow for 24 h prior to treatment. Cells were routinely observed and photographed by means of an IX70 inverted epifluorescence microscope (Olympus).

For 3D cultures, we seeded 80,000 cells mixed in 1:1 dilution of medium and Matrigel^®^ (Corning Inc.) in 24-well plates already coated with a solidified layer of pure Matrigel^®^. Cells were kept in these conditions for 72 h to allow microsphere formation prior to treatment. Cell-permeable DNA-binding dye Hoechst-33342 (Molecular Probes) was added (10 μg/mL final concentration in medium) for nuclei visualization.

### Chemicals and treatments

Dinaciclib (CAS 779353-01-4) was purchased from Selleckchem and diluted in DMSO (Sigma-Aldrich) to a 100 μM concentration. Used dilutions were made in complete medium. Pan-caspase inhibitor N-benzyloxycarbonyl-Val-Ala-Asp(O-Me) fluoromethyl ketone (z-VAD-fmk, CAS 187389-52-2) was purchased from ApexBio and diluted in DMSO to form a 50 mM stock solution. Used dilutions (40 μM) were made in complete medium. Generic BH3-mimetic ABT737 (CAS 852808-04-9) was obtained from ApexBio and diluted in DMSO to an initial 10 mM stock. The Bcl-x_L_ specific inhibitor A-1331852 (CAS 1430844-80-6) was purchased from ChemieTek and a 3 mM initial stock solution in DMSO was employed. The nucleoside analog Gemcitabine (CAS 122111-03-9) was obtained as hydrochloride salt from LC Laboratories and diluted in DMSO to form a 1 mM stock. In every case, further dilutions were made taking into account not to surpass 0.1% (v/v) content of DMSO in media.

### Proliferation and long-term clonogenic assay

For proliferation assessment, 3,000-5,000 cells were seeded in 96-well plates and treated with increasing concentrations of Dinaciclib, ABT737, A-1331852 or combinations of drugs for variable times up to 72 h. At desired time points, cells were incubated with WST-1 reagent (Roche) diluted 1:10 in complete medium up to 2 h in the incubator. Soluble formazan salt production from WST-1 was measured by absorbance (λ=440 nm) in a PowerWaveXS plate reader spectrophotometer (BioTek Instruments). IC_50_ values were calculated using Prism software (GraphPad).

Long-term analyses of post-treatment survival were performed by seeding 180,000-300,000cells in 6-well plates. Cells were cultivated with 25 nM Dinaciclib for 72 h prior drug wash-out and kept in culture up to a week. At that point cells were transferred to new plates to recover Dinaciclib Resistant strains or were stained with 0.5 % (w/v) Crystal Violet (Sigma-Aldrich) solution in 1:5 Methanol:PBS for 20 min and washed with water. Images reflect representative results of at least three independent experiments.

### siRNA experiments

Cells were transfected using DharmaFECT (Dharmacom, GE HealthCare) following manufacturer’s instructions. ON-TARGETplus Non-Targeting Control Pool (Dharmacom) was used as reference and customized siRNA sequences were purchased from Sigma-Aldrich:5’-AAUAACACCAGUACGGACGGG-3’ for Mcl-1 and 5’- ACAAGGAGAUGCAGGUAUUUU-3’ for Bcl-x_L_.

### Flow cytometry

For the simultaneous quantification of plasma membrane integrity and mitochondrial transmembrane potential (Δψ_m_), living cells were collected and stained with 1 μg/mL propidium iodide (PI, Molecular Probes) and 40 nM of DiOC_6_(3) (DiOC, Sigma-Aldrich) for 30 min at 37 °C. DiOC is a cyanine derivative that accumulates in functional mitochondria and dissipates when Δψ_m_ is lost. PI is a cell impermeable cationic stain (DNA intercalating) that accumulates in cell nuclei once the plasma membrane is no longer a barrier. So, we can identify cells committed to apoptosis (low levels of DiOC but still excluding PI), cells that suffered sudden necrosis (positive to PI but still retaining DiOC labeling) and cells that progress from apoptosis to secondary necrosis (negative to DiOC and positive to PI). Live cells (positive to DiOC and negative to PI) were not represented.

For cell cycle analysis, cells were fixed in 70 % ice-cold ethanol and labeled with 50 μg/mL PI in PBS containing 500 μg/mL RNase (Sigma-Aldrich). This approach also allows cell death quantification as subG_1_ events. Cytofluorometric determinations were performed employing a Gallios flow cytometer and data were statistically evaluated using Kaluza software (Beckman Coulter). Only the events characterized by normal forward scatter (FSC) and side scatter (SSC) parameters were included in sub-sequent analyses.

### Western blot analysis

Cells were lysed with RIPA Buffer (ThermoFisher Scientific) containing Protease Inhibitor Cocktail Tablets and Phospatase Inhibitor Cocktail Tablets (Roche). Lysates were sonicated and centrifuged at 13,000 rpm for 20 min (at 4 °C). Protein content was determined with BCA assay (Pierce Biotech.). Lysate aliquots (50 μg) were resolved by 8-12 % SDS-PAGE and transferred onto nitrocellulose membranes (Bio-Rad Laboratories). After blocking with 5 % skimmed milk in PBS containing 0.2 % (v/v) Tween-20 (Sigma-Aldrich) for 1 h (RT), membranes were incubated overnight at 4 °C with the appropriate primary antibody dilution (**Table 2**).

**Table 2:**
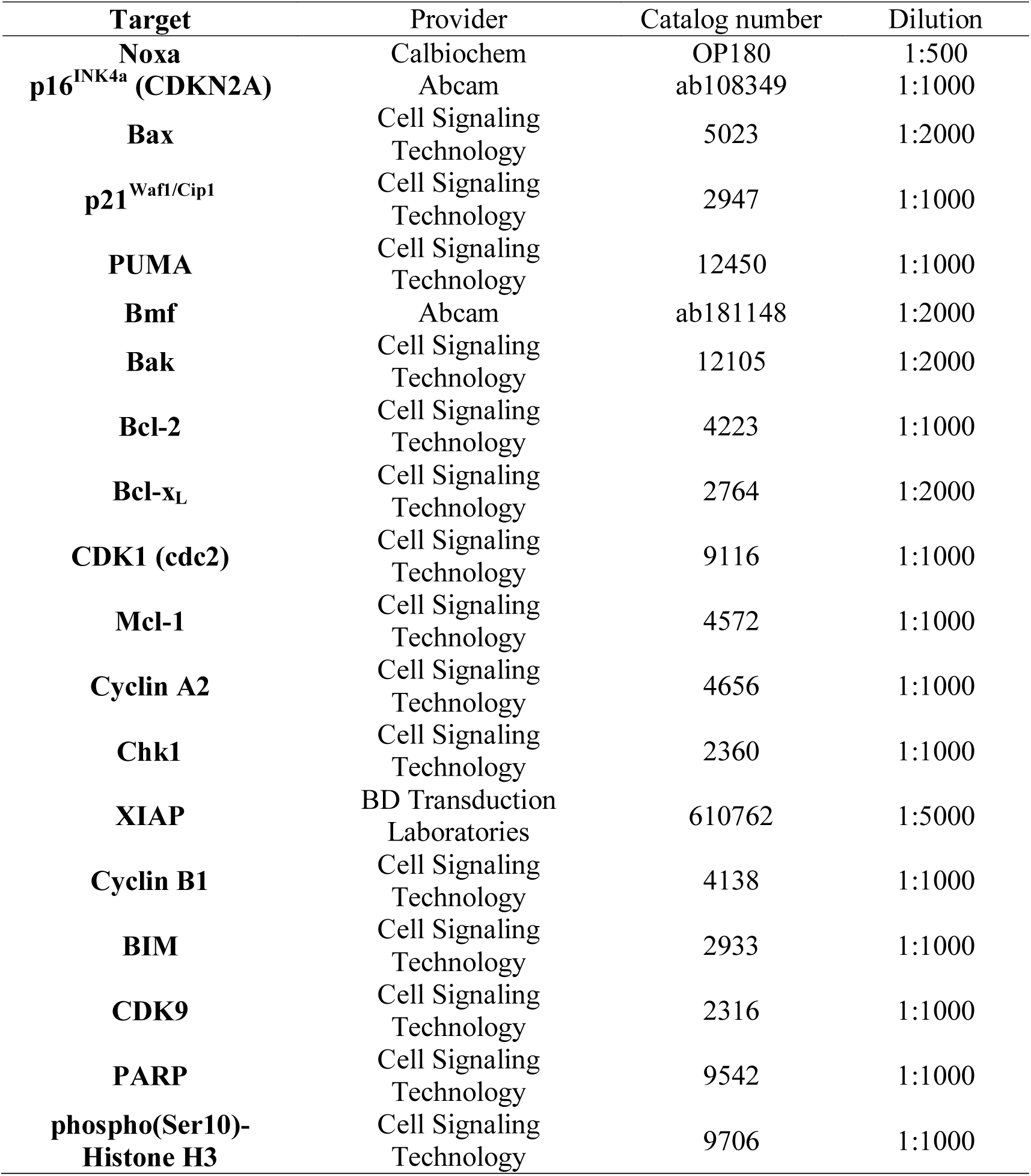
List of primary antibodies employed for western blotting and/or immunofluorescence.

Blots were then incubated at room temperature (RT) for 1 h with Goat anti-Rabbit/anti-Mouse IgG (H+L) horseradish peroxidase–conjugated secondary antibodies (#31460 and #31430, Thermo-Fisher Scientific) and bands were revealed in Hyperfilm ECL (Amersham) by enhanced chemiluminescence employing ECL Western Blotting Substrate (Pierce Biotech.). Immunodetection of β-actin (#ab49900) or α-tubulin (#ab28439, both from Abcam) were used as loading reference.

### Immunofluorescence

Cells (30,000-80,000) were seeded over glass coverslips placed into 24-well plates prior to treatment. At desired end-points, cells were fixed by 30 min incubation in 4 % (w/v) paraformaldehyde (Sigma-Aldrich) in PBS. Cells were sequentially permeabilized with 0.1 % (w/v) SDS in PBS (USB Corp.) and subjected to antigen blocking with 10 % (v/v) FBS in PBS (both for 20 min). Primary antibodies employed were Bax, p21^Waf1/Cip1^, p16^INK4a^ and phospho(Ser10)-Histone H3 (**Table 2**). Secondary antibodies were Goat anti-Mouse or anti-Rabbit IgG (H+L) Secondary Antibodies, conjugated either with Alexa Fluor^®^ 488 or 594 (ThermoFisher Scientific: #A-11001, #A-11005, #A-11008 and #A-11012). DNA counterstaining was performed upon 10 min incubation with 5 μg/mL solution of DAPI (Molecular Probes). Photographs were taken with either an Axio Observer.Z1 (Carl Zeiss AG) or a DM6000B (Leica Microsystems) epifluorescence microscope. Images were analyzed with Image J software (freely available from the National Institutes of Health at the address https://imagej.nih.gov/ij/).

### *In vivo* experimentation

For animal experimentation, Dinaciclib was purchased to ApexBio as powder and formulated in 20 % hydroxypropyl beta-cyclodextrin (HPβCD, Sigma-Aldrich) in deionized water. The formulation was stored at 5 °C, used within 7 days and warmed to RT and vortexed for 3 sec before intraperitoneal (i.p.) administration. ABT737 was also obtained from ApexBio and formulated in 30 % propylene glycol, 5 % Tween 80, 10 % DMSO, 3.3 % dextrose (all from Sigma-Aldrich) in water. Aliquots were kept at RT and vortexed before i.p. administration. A-1331852 was provided by Abbvie under Material Transfer Agreement and formulated in 60 % Phosal 50 PG (standardized phosphatidylcholine, PC, concentrate with at least 50 % PC and propylene glycol; gently provided by AbbVie), 27.5 % poly-ethylene glycol (PEG) 400, 10 % ethanol, and 2.5 % DMSO (all from Sigma-Aldrich). First, A-1331852 powder was suspended in DMSO and ethanol until a uniform cloudy suspension was obtained. PEG 400 and Phosal were then added and the solution was mixed by vortexing. The solution was allowed to sit for approximately 30 min after adding all the excipients helped to achieve a clear solution. A sonicator was also used for less than 10 min. Formulated compound was stored in an amber bottle at room temperature to protect it from light. A-1331852 formulation was administered *per os* (p.o.) by using an oral gavage. Dosages and administration regimes tested are summarized in **Supplementary Figure 5A & C**.

Female Athymic (Hsd:Athymic Nude*Foxn1*^*nu*^) mice (n=20) were purchased at Envigo. Female SCID^®^ Beige (CB17.Cg-*Prkdc*^*scid*^*Lyst*^*bg-J*^/Crl) mice (n=40) were acquired at Charles River. Animal care procedures were followed according to the Institutional Guidelines for the Care and Use of Laboratory Animals. Ethics approval was provided by the locally appointed ethics committee. IDIBELL animal facility abides to the Association for Assessment and Accreditation of Laboratory Animal Care (AAALAC) regulations.

Five million (5×10^6^) SK-LMS-1 cells in 1:1 RPMI:Matrigel^®^ were subcutaneously (s.c.) injected in the right dorsal flank of the mice. Tumor growth was followed by a caliper and volume was extrapolated using the formula vol=(L×l^2^)/2 were “L” refers to the major and “l” to the minor diameter measured. Drug treatments started when tumors reached a volume of approximately 100 mm^3^. Mice were blindly randomized into the different groups by using a 6-faced dice.

After the end of the experiment, tumors and organs of interest were extracted and fixed in 3.7 % Formol solution in PBS (Sigma-Aldrich) for 24 h. Afterwards, samples were paraffin embedded and prepared for Hematoxylin-Eosin (H&E) staining following standard protocols. Microscopical imaging was performed by means of a Nikon Eclipse 80i workstation. Photographs were analyzed using GNU Image Manipulation Program (GIMP) software (freely available at http://www.gimp.org).

### Statistical Analysis

Unless otherwise stated, experiments were performed thrice. Data were analyzed for statistical significance using Student’s *t* test, using either Calc (The Document Foundation) or Excel (Microsoft Corporation) software; *p* ≤ 0.05 was regarded as significant. Synergistic behavior of drugs in combination was assessed by means of the comparison of the surviving fractions according to the method of Valeriote & Lin (27). Mice survival curves were analyzed using Prism (GraphPad Software Inc.).

## Results

### Dinaciclib induces cell death in STSs independently of CDK1 and CDK9 expression levels

To have a base for inferring the response of STS to Dinaciclib, we analyzed the expression of CDK9, CDK1 and its partners Cyclin B1 and A2 in a series of cell lines representative of major groups of STS: synovial sarcoma SW982 and 1273-99, leiomyosarcoma SK-LMS-1 and SK-UT-1, myxoid liposarcoma 402-91 and 1765-92, dedifferentiated liposarcoma SW872 and myxofibrosarcoma HT-1080 (**Figure 1A**). Protein expression was variable among the cell lines, from the rather homogenous 43 kDa (low band) of CDK9 to the more variable CDK1. SW872 expressed comparatively undetectable levels of CDK1 and its partners. However, the levels of the 55 kDa (high band) of CDK9 were among the highest.

**Figure 1:**
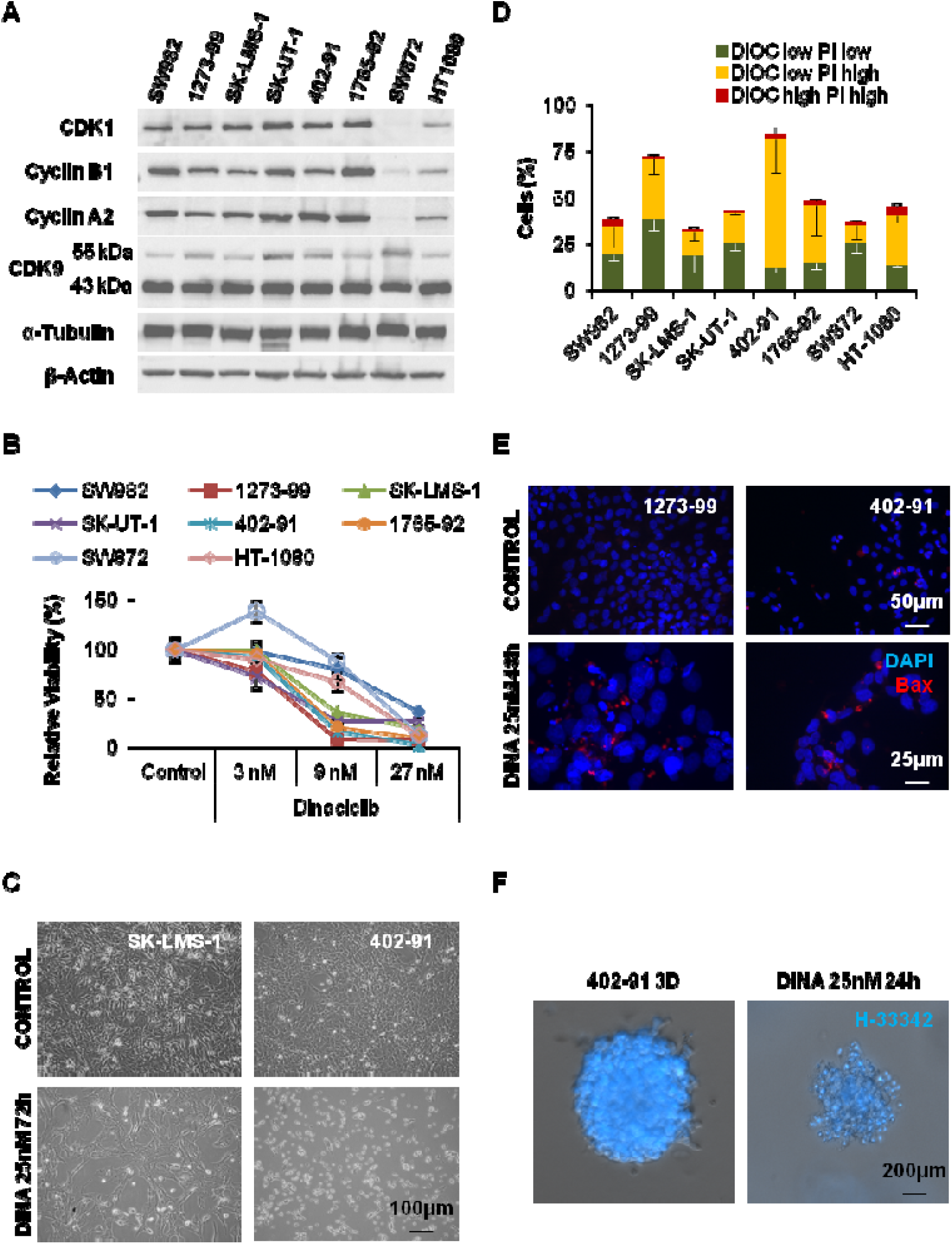
CDK inhibitor Dinaciclib is able to successfully inactivate sarcoma cell lines in a nanomolar range. (A) Representative western blot showing expression levels of Dinaciclib targets CDK1 and CDK9, and CDK1 partners Cyclin B1 and Cyclin A2 among different soft-tissue sarcoma derived cell lines. (B) Viability, measured by means of WST-1 reduction test, in a set of different STS cell lines incubated with increasing concentrations of Dinaciclib for 72 h; comparing with cells treated only with vehicle. (C) Microscopic imaging showing different cell behavior among cell lines when treated with Dinaciclib. (D) Comparison of apoptotic induction, cytofluorometric measurement of mitochondrial vital dye DiOC and death marker PI, in different STS derived cell lines treated with 25 nM Dinaciclib up to 72 h reveals different sensitivity among the cell lines. (E) Immunofluorescent detection of pro-apoptotic protein Bax showing its accumulation in *puncta* (mitochondria) in sensitive cells treated with 25 nM Dinaciclib. (F) 402-91 cells cultured as 3D spheroids by embedding into a Matrigel^®^ matrix and treated by Dinaciclib show loss of size and accumulation of pyknotic nuclei as revealed by staining with Hoechst-33342 (H-33342).

The same set of cell lines was used to test the ability of Dinaciclib to impair cellular viability (read as WST-1 assay, **Figure 1B**). SW982, SW872 and HT-1080 showed fewer decreases in viability at low concentrations, but at 27 nM differences among the cell lines were reduced. IC_50_ calculations failed for SW872 and HT-1080 due to high disparity of the data (**Supplementary Figure 1A**), but actual values should be close to the 12.760 nM obtained for SW982.

However, when observed under the microscope, the response to Dinaciclib was more diverse (**Figure 1C**). Rather than any form of cell death, most cell lines (like SK-LMS-1) showed a stop in proliferation with almost the same number of cells as seeded and few detached. Only 402-91 and 1273-99 showed an extensive accumulation of detached, dead cells. To clarify this dual effect, we performed an analysis of apoptotic induction by cytofluorometric quantification of the levels of the vital stains DiOC_6_(3) and propidium iodide (DiOC-PI). Comparison among the 8 cell lines treated with 25 nM Dinaciclib for 72 h evidenced the two major phenotypes from the microscopical observations (**Figure 1D**). Only 1273-99 and 402-91 presented values of apoptotic cell death exceeding the 50 %. Dinaciclib concentration of 25 nM was chosen under experimental conditions as the minimal value above which cell death reached a plateau as soon as 24 h of treatment either in responsive or in tolerant cell lines (**Supplementary Figure 1B**).

In order to confirm that Dinaciclib induced mitochondrial apoptosis we studied the translocation of pro-apoptotic protein Bax to the mitochondria (**Figure 1E**) and the caspase-3 activation reporter PARP (**Supplementary Figure 1C**). Both features were confirmed in the sensitive cell lines 1273-99 and 402-91. Furthermore, the addition of 40 μM pan-caspase inhibitor z-VAD-fmk to the system for 36 h was sufficient to abrogate the induction of cell death in both cell lines (**Supplementary Figures 1D and E**).

Dinaciclib potential as cell death inducer was conserved also in 3D tissue simulation. 402-91 cells embedded in a Matrigel^®^ matrix grew as microspheres. By treating the microspheres with Dinaciclib for 24 h prior staining with vital dye H-33342, we clearly observed a reduction of the size of the spheres but also the accumulation of bright pyknotic nuclei generated by apoptosis (**Figure 1F**).

Taken together, our results show the potential of Dinaciclib as inducer of apoptotic cell death in STS at nanomolar concentrations.

### Dinaciclib effects on cells last after wash-out

To complete our understanding of cellular response to Dinaciclib, we performed long-term experiments analyzing cell response after drug wash-out. During the first days after Dinaciclib removal, cell lines from the tolerant group (like SW982) underwent cell death albeit the identification of a number of viable surviving colonies (**Figure 2A**). Cells from the sensitive lines (like 1273-99) presented rare, if any, surviving colonies. Additionally, we allowed those colonies to grow into sub-cultures that we dubbed Dinaciclib Resistant (DR). Those DR lines were newly challenged with 72 h incubation of Dinaciclib 25 nM. DRs from the tolerant group did not show any evident difference in behavior from the parental culture. But those derived from the sensitive cell lines appeared to have acquired a better fitness against the drug (**Figure 2B**). Nevertheless, when different experiments were summed up there was no statistical significance in this trend to a reduced cell death induction (**Figure 2C**).

**Figure 2:**
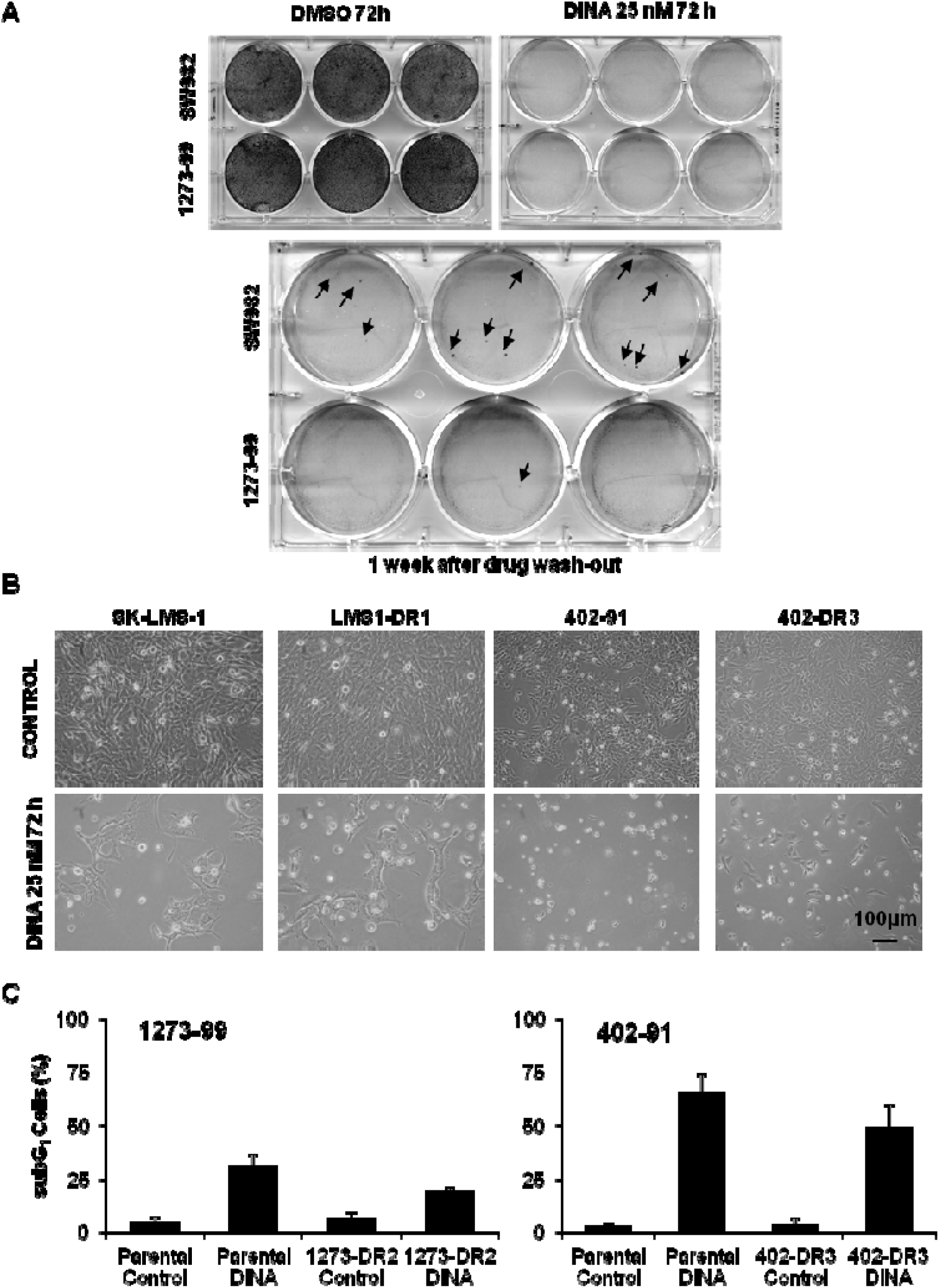
Dinaciclib did not drive strong selection of resistant clones after drug wash-out. (A) Crystal violet staining images showing resistant colonies (arrows) arising from highly sensitive 1273-99 cell line (low row) and more tolerant SW982 (upper row) after Dinaciclib wash-out. (B) Microscopic imaging showing polyclonal Dinaciclib Resistant (DR) strains response after being newly subjected to 72 h of treatment. (C) Cytofluorometric measurement of death induction (subG_1_ DNA content in PI profiles) for parental and derived DR strains from sensitive 1273-99 and 402-91 cell lines after a new cycle of 72 h incubation with Dinaciclib 25 nM (DINA).

Therefore, our data suggests that Dinaciclib exerts a long-term effect in cell viability with the additional value that surviving colonies do not show a strong selection towards drug resistance.

### Sensitive and tolerant cell lines differ in cell death protein expression patterns

As the known targets of Dinaciclib yielded no clear correlation with drug response (**Figure 1**), we performed an extensive analysis of proteins involved in cell cycle control and mitochondrial apoptosis among examples of responsive (402-91 and 1273-99) and tolerant (SW982 and SK-LMS-1) cell lines (**Figure 3A**).

**Figure 3:**
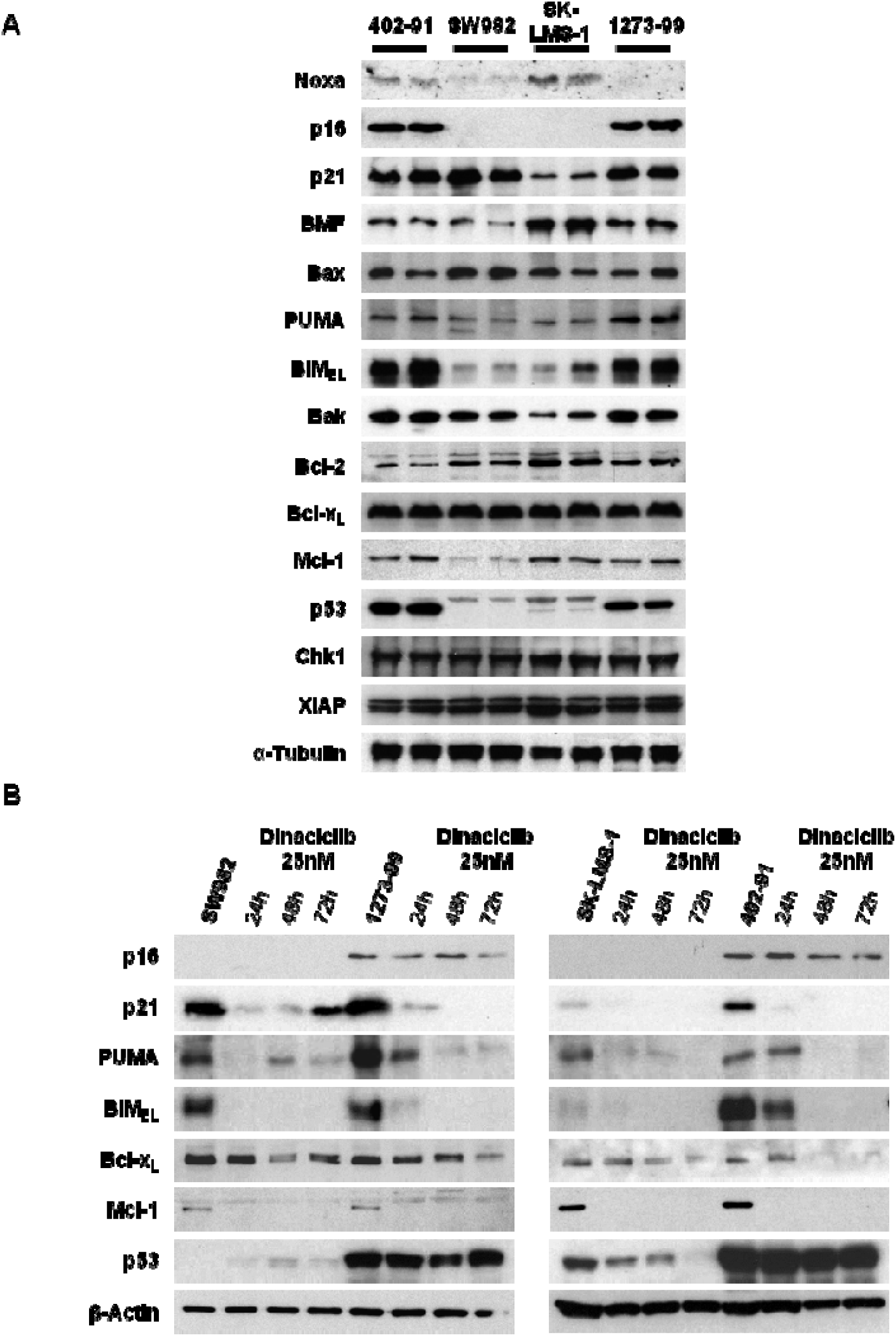
**Dinaciclib highly sensitive cell lines present characteristic levels of key regulators of the cell cycle and apoptosis triggering. (A) Representative western blots showing protein expression patterns in control duplicates from myxoid liposarcoma 402-91, synovial sarcoma 1273-99 and SW982 and leiomyosarcoma SK-LMS-1. A representative sample of equal charges marker** α**-tubulin is also showed. (B) Representative western blots showing the decay rhythm of key components of cell cycle and apoptosis regulation at selected end-points of Dinaciclib incubation. A representative sample of equal charges marker** β**-actin is also showed.**

For most proteins little variation among the cell lines was observed. However, other proteins did shown remarkable differences in expression between the two groups. Tumor suppressor p53 was almost undetectable in Dinaciclib tolerant cells when compared with the responsive ones. In this regard, it is important to note that SK-LMS-1 cell line has been described as *TP53* mutant (28). SK-LMS-1 expressed also relatively low levels of p21^Waf1/Cip1^ (p21) protein, a p53 target. Finally, CDK4 inhibitor p16^INK4a^ (p16) presented a pattern very similar to p53. If we compare this data with the study of cell cycle changes, we can see that only cell line SW982 presented a statistically significant arrest in G_2_/M phase (**Supplementary Figure 2A**). In any case, most of the Dinaciclib tolerant cell lines did share the same trend. To analyze the precise nature of this G_2_/M blockade we visualized treated and untreated cells immunolabelled for Ser10-phosphorylated Histone H3 (p-H3) and p21. Our results confirmed that the arrest was not due to accumulation of mitotic events but rather to the arrest of cells in the G_2_ phase (or in a tetraploid G_1_). In concordance with the protein expression data p21 was not induced in SK-LMS-1 cells (**Supplementary Figure 2B**). To complete the analysis, we found that senescence-related p16 translocated towards the nuclei in tolerant cell lines treated with Dinaciclib, in a way similar to the sensitive ones (whose signal was clearly more intense) (**Supplementary Figure 2C**). Thus, tolerance to Dinaciclib seems to be linked to the generation of a partial G_2_ arrest without a clear or sustained induction of senescence, as cells finally die in the following days (**Figure 2A**).

Among the mitochondrial apoptotic control machinery, major differences were found in the small pro-apoptotic BH3-only proteins PUMA and BIM. Both were found comparatively over-expressed in Dinaciclib sensitive cell lines. Minor differences were also found in Noxa, BMF or Mcl-1 but with no correlation with the Dinaciclib response.

The major effect of Dinaciclib via inhibition of CDK9 is the interruption of mRNA transcription (17,29). On this basis, we analyzed how our main targets evolved upon Dinaciclib incubation (**Figure 3B**). As expected, most of the proteins showed a radical decrease in expression after Dinaciclib incubation. In concordance with the data from **Supplementary Figure 2C**, p21 was clearly induced in SW982 cell line. Cell cycle modulators p16 and p53 levels were kept in Dinaciclib sensitive cells. More importantly, anti-apoptotic regulator Bcl-x_L_ expression seemed invariable or slightly reduced by Dinaciclib. Only in 402-91 cells repression of Bcl-x_L_ was observed after 48 h of treatment. In contrast, Mcl-1 was early and completely suppressed in all cell lines. Finally, Dinaciclib responsive cell lines retained for longer time the, already higher, levels of pro-apoptotic BIM and PUMA.

Summing up, differences in key proteins among STS cell lines can explain particular features of Dinaciclib response.

### Bcl-x_L_ silencing enhances sensitivity to Dinaciclib in SW982 and SK-LMS-1 cell lines

Based on our protein expression data, we developed a hypothetical framework to study the mechanistic of Dinaciclib action. Dinaciclib sensitive cells undergo quick apoptosis triggering due to their relative retention of higher levels of pro-apoptotic proteins (BIM, PUMA) after quick disappearing of anti-apoptotic Mcl-1. The high levels of p16 prevent surviving cells to get arrested in G_2_ phase (**Figure 4A**). On the other hand, Dinaciclib tolerant cells had relatively lower levels of BIM and PUMA, and they are lost faster after CDK9 inhibition than in the responsive ones. Remaining Bcl-x_L_ prevents apoptosis to be triggered and cells proceed to G_2_ when other mechanisms (like p21 induction in SW982) are activated (**Figure 4B**). In time, the induced damage will force the cells to die (**Figure 2A**).

**Figure 4:**
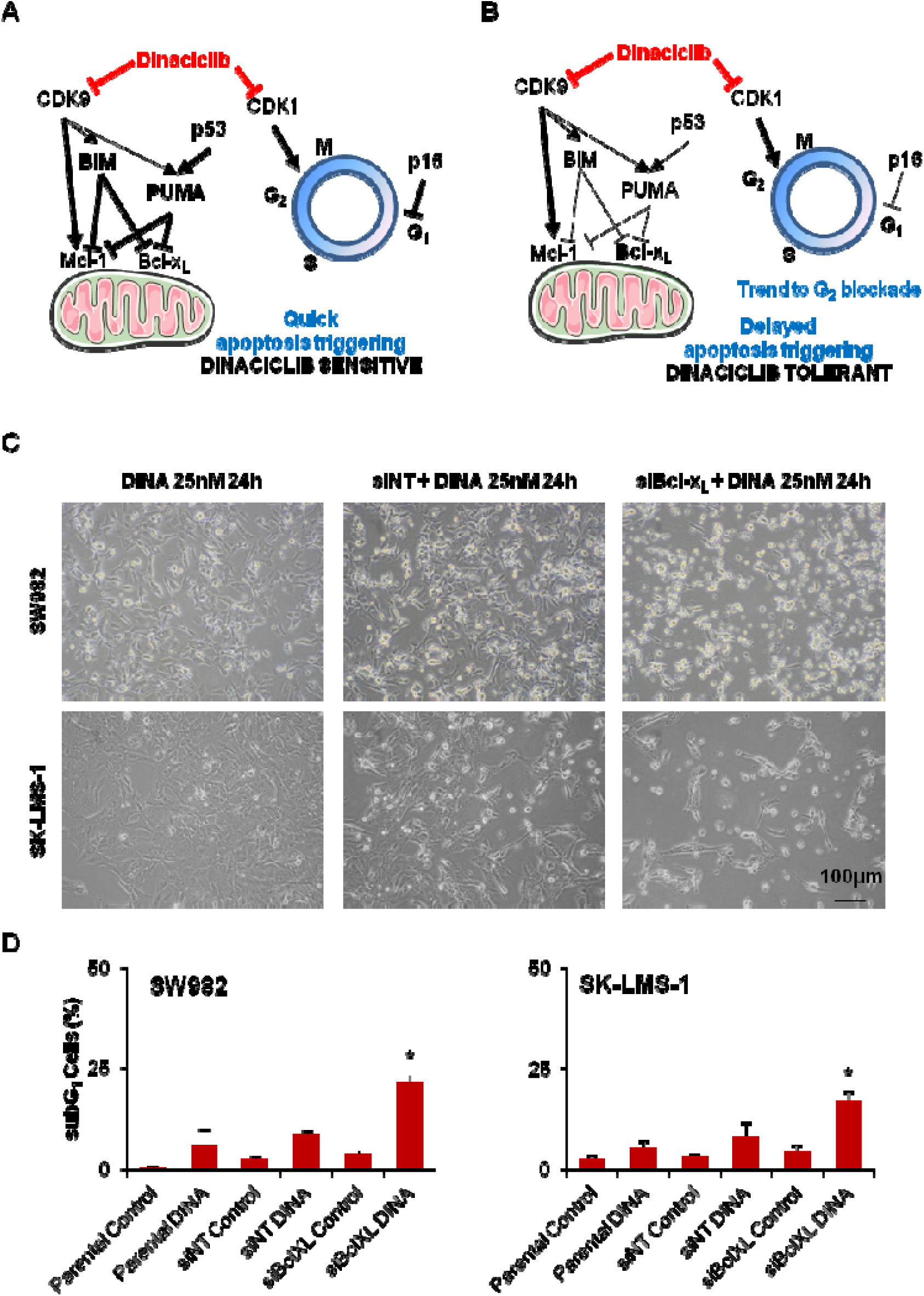
Bcl-x_L_ is the key regulator of apoptosis triggering after Dinaciclib treatment. (A & B) Schematics of the proposed mechanisms of action for the triggering of apoptosis in STS cell lines sensitive (A) and tolerant (B) to Dinaciclib. (C) Representative microscopic imaging of SW982 (above) and SK-LMS-1 (below) cell lines treated with 25 nM Dinaciclib for 24 h without prior challenge or after 24 h treatment with alternatively a non targeting (siNT) or a Bcl-x_L_ directed (siBcl-x_L_) siRNA. (D) Cytofluorometric quantification of cell death induction (measured as subG_1_ DNA content in PI profiles of fixed cell populations) after sequential treatment with siRNA and Dinaciclib. Data are presented as means ± SD. Statistical significance was achieved by the Student’s *t* test from at least three different experiments: \**p* ≤ **0.05.**

To check the coherence of our hypothesis, we have reviewed the data presented by Booher and co-workers in 2014 showing correlation between *MCL1:BCL2L1* mRNA ratio and viability after 24 h 100 nM Dinaciclib treatment (18). By reducing the 250 samples to the ones appertaining to STS, we still found a statistically significant correlation, meaning that those cell lines more depending on Mcl-1 were also prone to die after Dinaciclib (**Supplementary Figure 3A**).

For experimental validation, we addressed the role of Mcl-1 in Dinaciclib-induced cell death by siRNA silencing (**Supplementary Figure 3B**). Although Mcl-1 was successfully depleted, no major increase in cell death was observed in SW982 cells (**Supplementary Figure 3C**). However, depleting Bcl-x_L_ in Dinaciclib tolerant cell lines before Dinaciclib addition (**Supplementary Figure 3D**) clearly resulted in an increase of the detachment of cells (**Figure 4C**). When cell death induction was quantified, the increase of cell death at 24 h of treatment was deemed statistically significant (**Figure 4D**).

Therefore, our results supported our hypothesis that Bcl-x_L_ is the more relevant counteracting mediator of Dinaciclib-induced apoptosis in STS.

### BH3-mimetics are able to synergistically enhance and accelerate cell death in STS

To assess the possibility of a combination therapy to enhance Dinaciclib action we chose two different BH3-mimetics: one with a broad action spectrum, but excluding Mcl-1 (ABT-737) and one specific only for Bcl-x_L_ (A-1331852). BH3-mimetics activity over Dinaciclib tolerant cells as monotherapy was negligible (**Supplementary Figure 4A & C**) but its effect was notable when combined with 25 nM Dinaciclib (**Supplementary Figure 4B & D**). Cytofluorometric measurement showed that both combinations synergized (27) in terms of apoptosis induction after 24 h of treatment at very low concentrations (**Figure 5A-B**). Moreover, A-1331852 was more efficient than ABT-737. These results confirmed our operational hypothesis regarding Dinaciclib mechanism of action.

**Figure 5:**
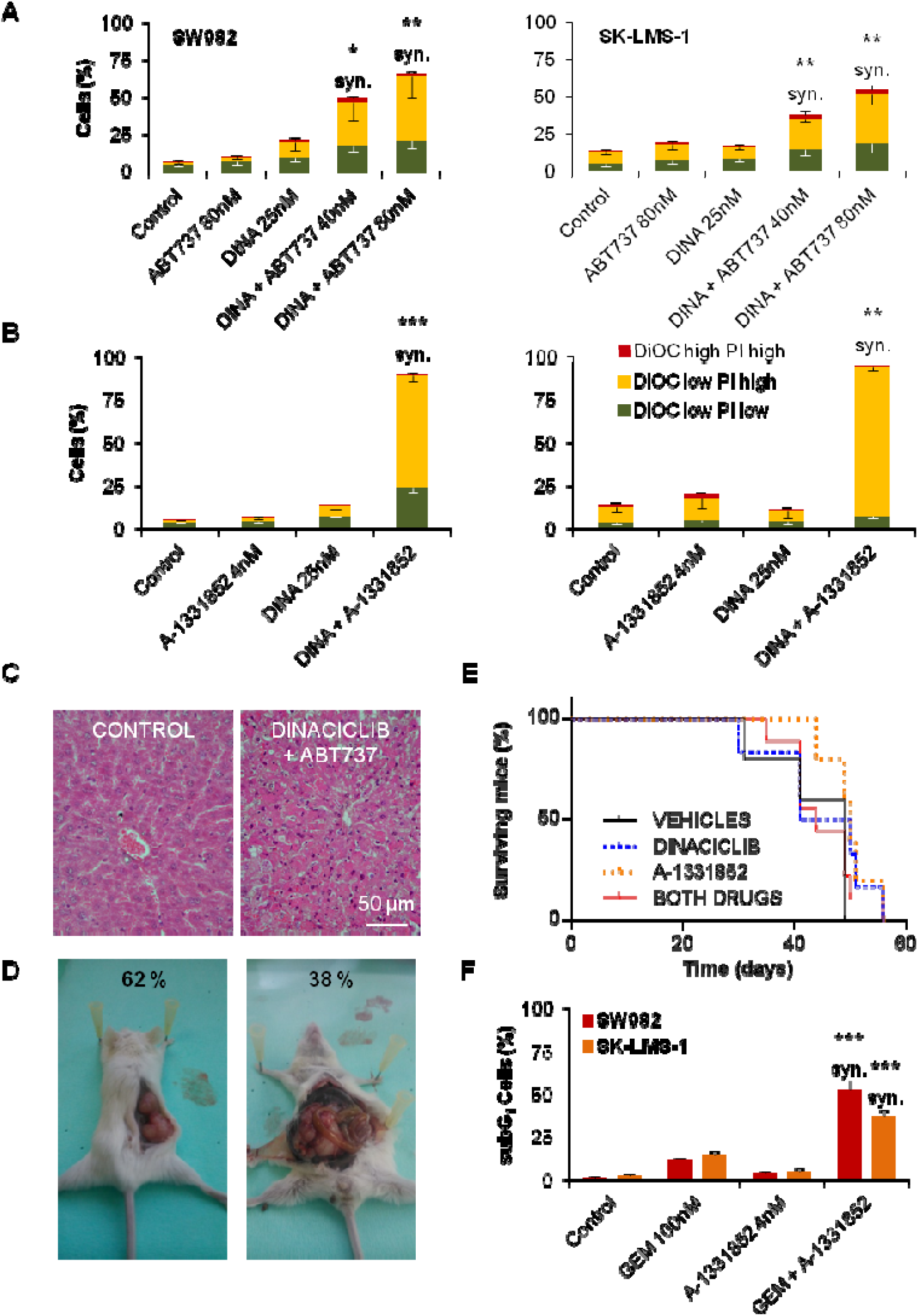
BH3-mimetics targeting Bcl-xL synergize with Dinaciclib to achieve quick apoptosis. (A) Cytofluorometric quantification of cell death via DiOC-PI staining of SW982 (left) and SK-LMS-1 (right) cell lines treated for 24 h with Dinaciclib 25 nM, ABT-737 80 nM or a combination of both drugs. (B) Cytofluorometric quantification of cell death via DiOC-PI staining of SW982 (left) and SK-LMS-1 (right) cell lines treated for 24 h with Dinaciclib 25 nM, A-1331852 4 nM or a combination of both drugs. (C) Microscopical imaging of H&E stained liver sections from Hsd:Athymic Nude*Foxn1*^*nu*^ mice control (left) and treated (right) after 3 i.p. injections with 40 mg/Kg Dinaciclib and 7 i.p. injections of ABT-737 who suffered sudden death at day 8. (D) Photographs showing typical subcutaneous tumor (left) and atypical intra-peritoneal tumor (right) caused by SK-LMS-1 cells in CB17.Cg-*Prkdc*^*scid*^*Lyst*^*bg-J*^/Crl mice. Percentages of occurrence are presented on the images. (E) Kaplan-Meier graph showing the survival of CB17.Cg-*Prkdc*^*scid*^*Lyst*^*bg-J*^/Crl mice inoculated (s.c.) with 5×10^6^ SK-LMS-1 cells and afterwards treated with i.p. 20 mg/Kg Dinaciclib, p.o. 25 mg/Kg A-1331852 or both drugs. Control mice were treated with both drugs vehicles. Time 0 represents the beginning of the treatment. (F) Cytofluorometric quantification of cell death (as subG_1_ in PI profiles in fixed cells) of SW982 and SK-LMS-1 cells treated with 100 nM Gemcitabine (GEM), 4 nM A-1331852 or a combination of both drugs for 48 h. Data are presented as means ± SD. Statistical significance was achieved by the Student’s *t* test from at least three different experiments: \**p* ≤ **0.05; \*\**p*** ≤ **0.001;\*\*\**p*** ≤ **0.0001. syn. denotes synergistic behavior as determined according Valeriote & Lin.**

The next logical step was trying the drug combination *in vivo*. Our first approaches were aimed to assess treatment toxicity and cell growth using Hsd:Athymic Nude*Foxn1*^*nu*^ mice. We tried different combination regimes of Dinaciclib and ABT-737 based on literature (18,30) (**Supplementary Figure 5A**), but we always observed liver toxicity linked to massive hepatocyte death (**Figure 5C**). Moreover, human SK-LMS-1 showed a poor efficiency of nesting and growing as sub-cutaneous tumors in those mice (**Supplementary Figure 5B**). In consequence, we moved to the more immunodepressed CB17.Cg-*Prkdc*^*scid*^*Lyst*^*bg-J*^/Crl mice trying a lower dosage of Dinaciclib (as it was able to kill the mice even as monotherapy) and A-1331852 instead of ABT-737 (**Supplementary Figure 5C**). SK-LMS-1 cells grew better in these mice **(Supplementary Figure 5D**), but an important percentage of tumors appeared as intra-peritoneal masses despite being injected sub-cutaneously (**Figure 5D**). Intra-peritoneal tumors grew by surrounding, engulfing and finally invading the kidney (**Supplementary Figure 5E**). Those mice were excluded from the final experiments. Final *in vivo* testing of anti-tumoral combined activity of both drugs was unsuccessful in terms of mice survival (**Figure 5E**). The failure was allegedly due to a lack of overall effect of the drugs, as can be inferred for the unchanged mice weight (**Supplementary Figure 5F**). Summing up, we could not reach a therapeutic regime in which the drugs showed anti-tumor activity without induction of deadly toxicity.

Notwithstanding this set back, we sought to understand whether the key role of Bcl-x_L_ in Dinaciclib-induced cell death was a particular feature or part of a more general behavior. Other groups have already shown a positive cooperation of BH3-mimetics with CDK inhibitors and anthracyclines (26,31). So, we tested another commonly used anti-tumor drug in STS therapy: nucleoside analogue Gemcitabine. Our results showed that A-1331852 synergized with Gemcitabine in terms of cell death induction after 48 h of combined treatment (**Figure 5F**).

Taking together, our results unveil the specificities of cellular response to CDK inhibitor Dinaciclib and positively identify Bcl-x_L_ as the key player of apoptosis induction in these systems. More pharmaceutical research will be required to achieve a safe way for clinical practice, probably through the development of new delivery methods or new molecules with reduced side-effects. Still, this question falls beyond the scope of our study.

## Discussion

Dinaciclib is one of the most promising compounds in the group of CDK inhibitors because is able to act by meddling in the progression of late steps of the cell cycle, the entry in mitosis, transcriptional regulation and even the unfolded protein response (15,32,33). It has been studied in clinical trials on cancer entities such as acute leukemia and other pediatric malignancies (11,14,20). Dinaciclib was well tolerated, but its therapeutic effect as single agent did not achieve any total remission. However, in several neoplasias it is considered a good candidate for combination therapies (7,33,34), including BH3-mimetics (26).

Here we present new data on Dinaciclib pre-clinical results in STS cell lines. Our results show that Dinaciclib is able, as single agent, to impede cell proliferation in a nanomolar range (**Figure 1B**) and to induce cell death at least in myxoid liposarcoma 402-91 and synovial sarcoma 1273-99 cells (**Figure 1D**). Our model of 3D culture also proves that Dinaciclib is able to infiltrate and kill cells in solid tumors (**Figure 1F**). These results align with previous pre-clinical reports that have also proved the utility of Dinaciclib in different tumor entities (16,18,33). Moreover, our long-term studies after Dinaciclib removal showed that the deregulation induced by the drug was enough to promote a delayed death (five days after drug withdrawal) and that the remnant colonies did not develop an enhanced resistance to re-challenge (**Figure 2C**). The lack of reinforced tolerance to Dinaciclib in DR lines opens the gate to the establishment of repeated administrations or combination procedures (35,36).

To do so, a proper cell death mechanism characterization is paramount (37). In this sense, our results proved that the main mechanism involved in Dinaciclib killing is the mitochondrial apoptotic pathway (**Figure 1D-E and Supplementary Figure 1B-E**). Other *in vitro* studies published with Dinaciclib also pointed to apoptotic mechanisms but they provided a less detailed characterization of the cell death (16,17,19). Dinaciclib action involves two clearly different processes: cell cycle inhibition and alteration in protein production and homeostasis (9,32). The study of those cell lines more tolerant to Dinaciclib is helpful as their delayed death allow us to observe other variables. Cell cycle profiling showed a trend to an arrest in G_2_-to-M transition that was only significant in SW982 (**Supplementary Figure 2A**). SW982 cells are also the only cell line able to induce an increase in p21 protein (**Figure 3B and Supplementary Figure 2B**). Our data agreed to previous reports in other cancer entities, in which the cell cycle arrest was revealed when apoptosis was inhibited by Bax/Bak DKO (18). But this mechanism may not be universal as most of our Dinaciclib tolerant cell lines did not reach statistically significant mitotic arrest. However, the sensitive cell lines indeed did increase their G_2_/M and postM (tetraploidy) phases when apoptosis was abrogated by z-VAD-fmk (data not shown). So, it appears that the effect on the cell cycle deregulation in our models may be an important contributor in the apoptotic triggering. Further research is needed to understand if other proteins (like p21) are required for G_2_ arrest after CDK1 inhibition.

The role of cell cycle regulator p16 is intriguing. It is one of the multiple transcripts of the *CDKN2A* locus, all of them with key functions in the homeostasis of the cell cycle, tumor suppression, senescence and aging (38–42). The expression of p16 is not directly regulated by p53 but rather by Jun-B and Ras via Ets-1/2 (38). However, there are references cross-linking Ets-1/2 transcription factors with Bcl-2 family expression (43–46) and even p53 activity (47). In our settings, the sensitive cell lines 1273-99 and 402-91 exhibit higher levels of p16 than the tolerant ones (**Figure 3A and Supplementary Figure 2C**). Hence, there is supporting data to consider that expression levels of p16 could be linked to an apoptotic-permissive unbalance among the Bcl-2 family that enhances Dinaciclib sensitivity. As has been proved in other cancer entities, we consider interesting to explore the use of p16 as a predictive marker for cell cycle targeted therapies (39,42).

Dinaciclib causes a general depletion of protein content in the cell via RNA Polymerase II dephosphorylation with different impact depending on protein distinctive half-life (7,9,17,18). From our list of proteins of interest (**Figure 3B**), we focused in Bcl-x_L_ as it maintained its levels essentially constant after Dinaciclib incubation. Delayed depletion of Bcl-x_L_ was seen only in the most sensitive cell line (402-91). In contrast, PUMA and BIM levels decreased later in Dinaciclib responsive cell lines. Mitochondrial apoptosis is basically regulated by the inhibition status of large anti-apoptotic members of Bcl-2 family of proteins. This inhibition status relies on the balance with the pro-apoptotic members of the group, particularly the small BH3-only proteins (48). Published literature has focused in anti-apoptotic regulator Mcl-1 levels as the main mediator of Dinaciclib cytotoxicity because its expression drops drastically after treatment (17–19). Recently, a more comprehensive study from Inoue-Yamauchi and co-workers lead to a more complex panorama in which every member of the Bcl-2 family has a role in the apoptotic triggering (26). For STS, the limited cohort from Booher and co-workers provides correlation between Dinaciclib sensitivity and Mcl-1/Bcl-x_L_ ratio (**Supplementary Figure 3A**). However, our data showed that Mcl-1 levels are already low in SW982 cells (**Figure 3A**) and quickly become almost undetectable upon Dinaciclib treatment in all cell lines (**Figure 3B**). So, we envisaged a mechanistic hypothesis that put remnant Bcl-x_L_ interaction with BIM and/or PUMA in the decision point of apoptosis triggering in STS cell lines (**Figure 4A-B**). When Dinaciclib is added, its effect over CDK9 causes a quick depletion in anti-apoptotic protein Mcl-1 but the effect over BIM or PUMA is weaker and negligible on p53. PUMA and BIM are thus capable to initiate mitochondrial apoptosis by inhibition of Bcl-x_L_. Yet SW982 and SK-LMS-1 have low levels of BIM and PUMA. In the presence of Dinaciclib, the remaining levels of Bcl-x_L_ are enough to sustain cell viability for longer time. Then, the effects over the cell cycle (via CDK1) become detectable as a trend (significant in SW982) to a G_2_ blockade. Silencing experiments with siRNA confirmed our suspicion, as Bcl-x_L_ induced a clear increase in detached cells whereas Mcl-1 inhibition exerted no major difference (**Figure 4C-D and Supplementary Figure 3B-C**).

Bcl-x_L_ involvement as key mediator of apoptosis triggering has been previously described in several contexts including cell cycle alterations and other sarcoma entities (49–51). Among the different Bcl-2 family interactions, Bcl-x_L_ has strong affinity for pro-apoptotic BH3-only proteins PUMA and BIM (49,50,52,53). BH3-mimetics are a new class of drugs precisely designed to benefit of this interaction to promote cell death (24). Since the recent FDA approval of Venetoclax, these molecules are in the spotlight of cancer research, albeit some groups are calling to improve the accurateness of data analysis (54). A more detailed research in their precise mechanism of action is needed in order to design meaningful combinations for treatment. Nevertheless, we decided to go ahead and test our working hypothesis by designing drug combination testing *in vitro*. Two BH3-mimetics were chosen: ABT-737, a rather generic compound that, inhibits Bcl-2, Bcl-w and Bcl-x_L_ proteins (55) and A-1331852, a newer and Bcl-x_L_ specific molecule (56). Neither of them induced a relevant cell death accumulation as monotherapy, but when combined with Dinaciclib they showed a fast synergistic induction of cell death in both SW982 and SK-LMS-1 cells (**Figure 5A-B and Supplementary Figure 4**). The effects were clearly better with A-1331852 than with ABT-737, providing more support to our mechanistic hypothesis. Regretfully, we encountered too many problems at escalating our settings to *in vivo* experimentation (**Figure 5C-E and Supplementary Figure 5**). The main issue was the development of deadly liver toxicity, that was already glimpsed in literature (57). Our data supports the need of more pharmaceutical research in order to generate a safer treatment protocol.

In the past few years, other groups have recognized the potential of BH3-mimetics in combination approaches to kill cancer cells. As previously commented, Inoue-Yamauchi and co-workers, using screening technologies in a small-cell lung cancer model, found that anti-tumor properties of CDK9 inhibitors and anthracyclines were boosted when combined with those molecules (26). Research in STS has also pointed to the suitability of such combinations. Possible combinations for BH3-mimetics include anthracyclines, histone deacetylase inhibitors, tubulin poisons and etoposide (31,58–60). Here, we add to the list Dinaciclib and Gemcitabine (**Figure 5F**) and we point that the major focus should be put in Bcl-x_L_ inhibitors. Both, our data and the previous literature support the conclusion that Bcl-x_L_ targeted molecules behave better than generic BH3-mimetics. The existence of a single molecular route to explain this multiple activity seems unlikely. Bcl-x_L_ has been identified as a key player of cell death triggering after mitotic arrest (50). This can explain the synergy reported with tubulin poisons or other compounds that induced strong mitotic arrest. But, it is not the case with Dinaciclib, as we obtained no relevant mitotic arrest linked to the cell death triggering. The G_2_ blockage reported for SW982 occurs at 72 h and combined cell death is verified at 24 h. Thus, in our setting it is more relevant the quick loss of protein players once Dinaciclib has disturbed the process of protein synthesis. The higher life time of Bcl-x_L_ positions it as the cornerstone of apoptotic induction. For Gemcitabine we cannot discard that the mechanism proceeds more similarly to the observed for mitotic disruptors. In this case, combined action takes more time to be verified (48 h *vs.* 24 h with Dinaciclib) showing that BH3-mimetics did not exert a pro-apoptotic activity by themselves. Indeed, their function is simply to push towards cell death *after* another injury has been detected in the cell.

Taking together, we have extensively showed that Dinaciclib (or an improved CDK9 inhibitor) is a suitable candidate for clinical trials in STS. It is capable to effectively kill STS cell lines in a relevant fashion, without inducing a strong selection towards drug resistance. Moreover, when combined with BH3-mimetics its action gets boosted in a remarkable way. Pharmacological research is, however, required in order to address the elevated toxicity linked to this drug.

## Acknowledgments

The mitochondria draw in Figure 4, was taken from Servier Medical Art (https://smart.servier.com/) under the terms of a Creative Commons Attribution 3.0 Unported License. The mice draw in Supplementary Figure 5 was taken and adapted from Wikimedia Commons image archive(https://commons.wikimedia.org/wiki/File:Mouse_line_drawing.jpg), licensed under the Creative Commons Attribution 2.5 Generic license. We thank David Herrero-Martín for helpful discussion.

## Author’s Contributions

**Conception and design:** S. R-V., O. M. T. and X. GdM. **Development of methodology:** S. R-V., M. F-G. and O. M. T. **Acquisition of data:** S. R-V., M. F-G., R. L-A., A. C-P., N. M-M. and S. G-M. **Analysis and interpretation of data:** S. R-V., M. F-G. and O. M. T. **Writing and/or revision of manuscript:** S. R-V., O. M. T. and X. GdM. **Funding acquisition:** S. R-V., O. M. T. and X. GdM.

---

**Figure S1:**
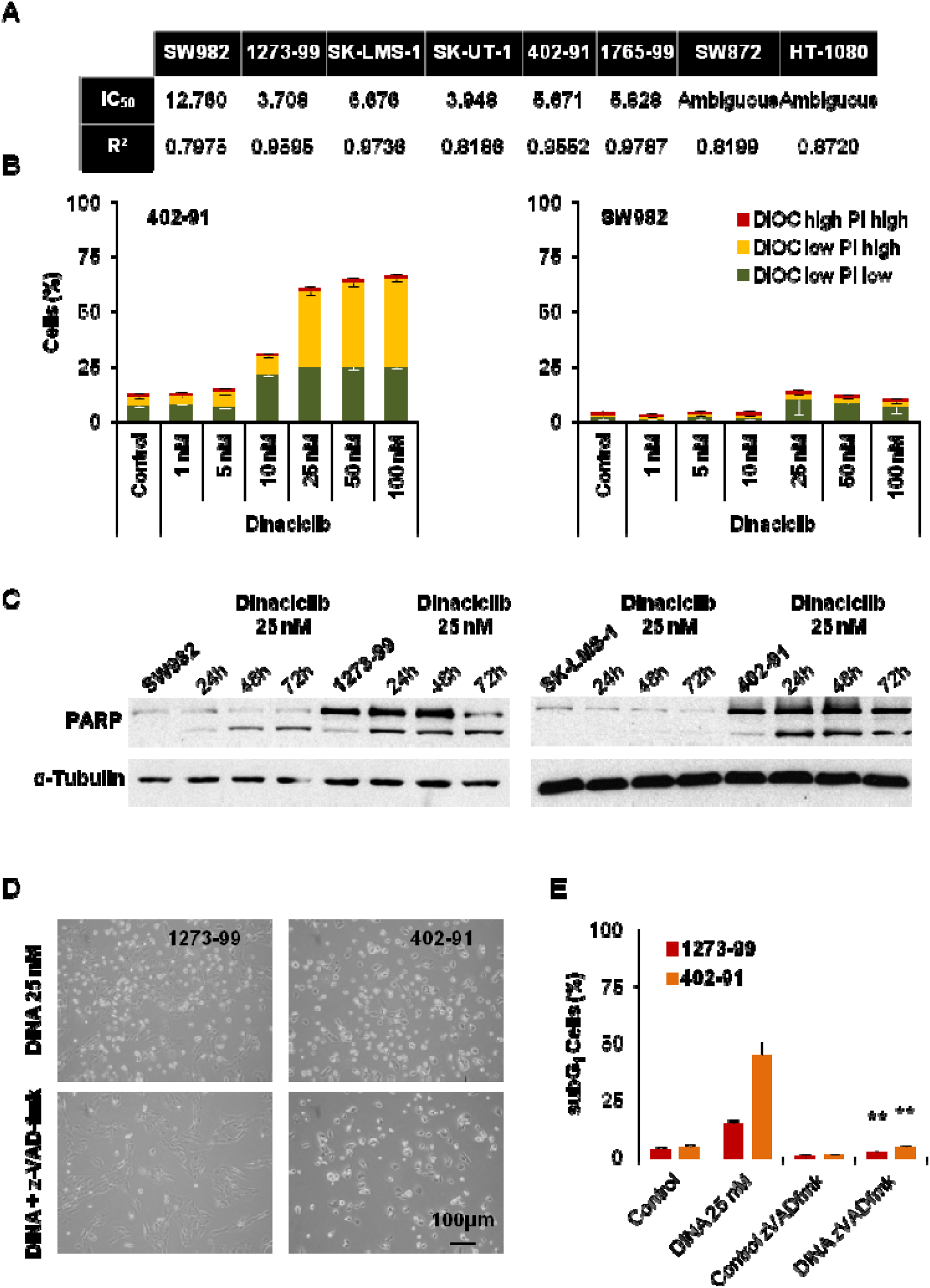
(related to Figure 1)**: Dinaciclib successfully induce apoptosis in sensitive cell lines.** (A) IC_50_ values obtained from WST-1 reduction test represented in Figure 1B. (B) Cytofluorometric measurement of mitochondrial vital dye DiOC and death marker PI showing apoptotic induction in 402-91 cell line (left), but not in SW982 cells (right), when treated with increasing concentrations of Dinaciclib for 24 h. (C) Representative western blots showing PARP protein cleavage after treatment with 25 nM Dinaciclib at different end times. (D) Representative microscopic imaging showing that Dinaciclib-induced apoptosis can be abrogated with co-incubation with 40 μM z-VAD-fmk for 36 h. (E) Cytofluorometric quantification of cell death (as subG_1_ in PI profiles in fixed cells) of 1273-99 and 402-91 cells treated with 25 nM Dinaciclib and 40 μM z-VAD-fmk for 36 h. Data are presented as means ± SD. Statistical significance was achieved by the Student’s *t* test from at least three different experiments: \*\**p* ≤ 0.001.

**Figure S2:**
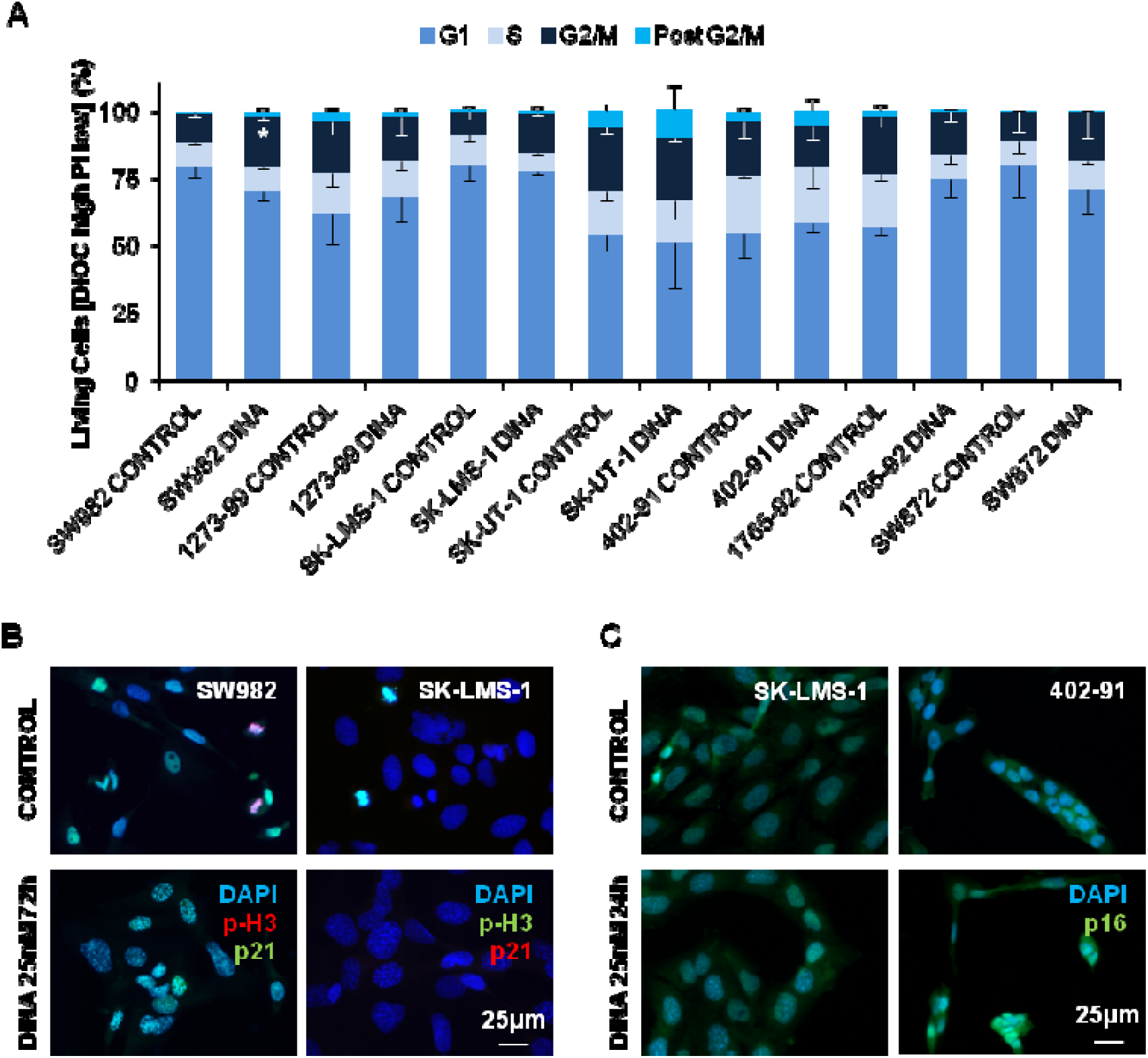
(related to Figure 3)**: Differences in protein expression levels explain particular responses to Dinaciclib treatment.** (A) Cytofluorometric measurement of cell cycle phases by means of H-333258 profiling of DiOC high PI low (living) cells in control conditions or after 72 h treatment with 25 nM Dinaciclib (DINA). SubG_1_ cells were excluded. (B) Microscopic imaging showing mitotic marker phosphorylated histone H3 (p-H3) and checkpoint protein p21^Waf1/Cip1^ (p21) in control conditions or after 72 h treatment with 25 nM DINA. (C) Microscopic imaging showing senescence marker p16^INK4a^ (p16) in control conditions or after 24 h treatment with 25 nM DINA. Data are presented as means ± SD. Statistical significance was achieved by the Student’s *t* test from at least three different experiments: \**p* ≤ 0.05.

**Figure S3:**
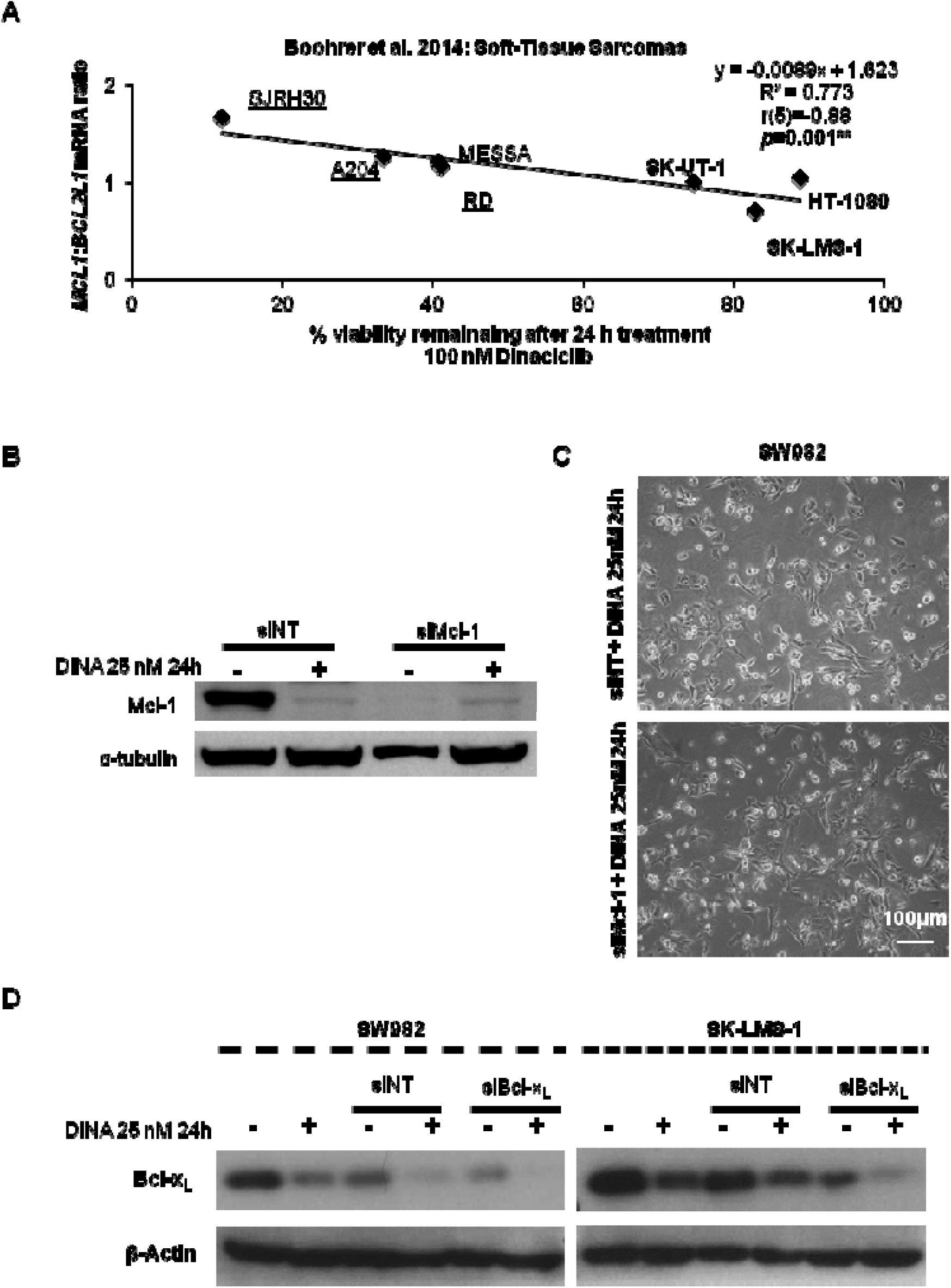
(related to Figure 4)**: Mcl-1 is not involved in Dinaciclib tolerance in STS cell lines.** (A) Reanalysis of Boohrer *et al.* data on Dinaciclib tolerance and Mcl-1 and Bcl-x_L_ expression (mRNA) constrained to the “Soft-Tissue Sarcoma” subset. Cell lines names are noted. Rhabdomyosarcoma cell lines are underlined. Cell lines also included in our study are highlighted in bold. (B) Representative western blot showing Mcl-1 expression in SW982 cell line in the combined presence of siRNA constructs and Dinaciclib. (C) Representative microscopic imaging showing SW982 cell response to sequential silencing of Mcl-1 prior treatment with Dinaciclib 25 nM. (D) Representative western blot showing Bcl-x_L_ expression in SW982 and SK-LMS-1 in sequential silencing of the target prior of Dinaciclib incubation. Data are presented as means ± SD. Statistical significance was achieved by the Student’s *t* test from at least three different experiments: \*\**p* ≤ 0.001.

**Figure S4:**
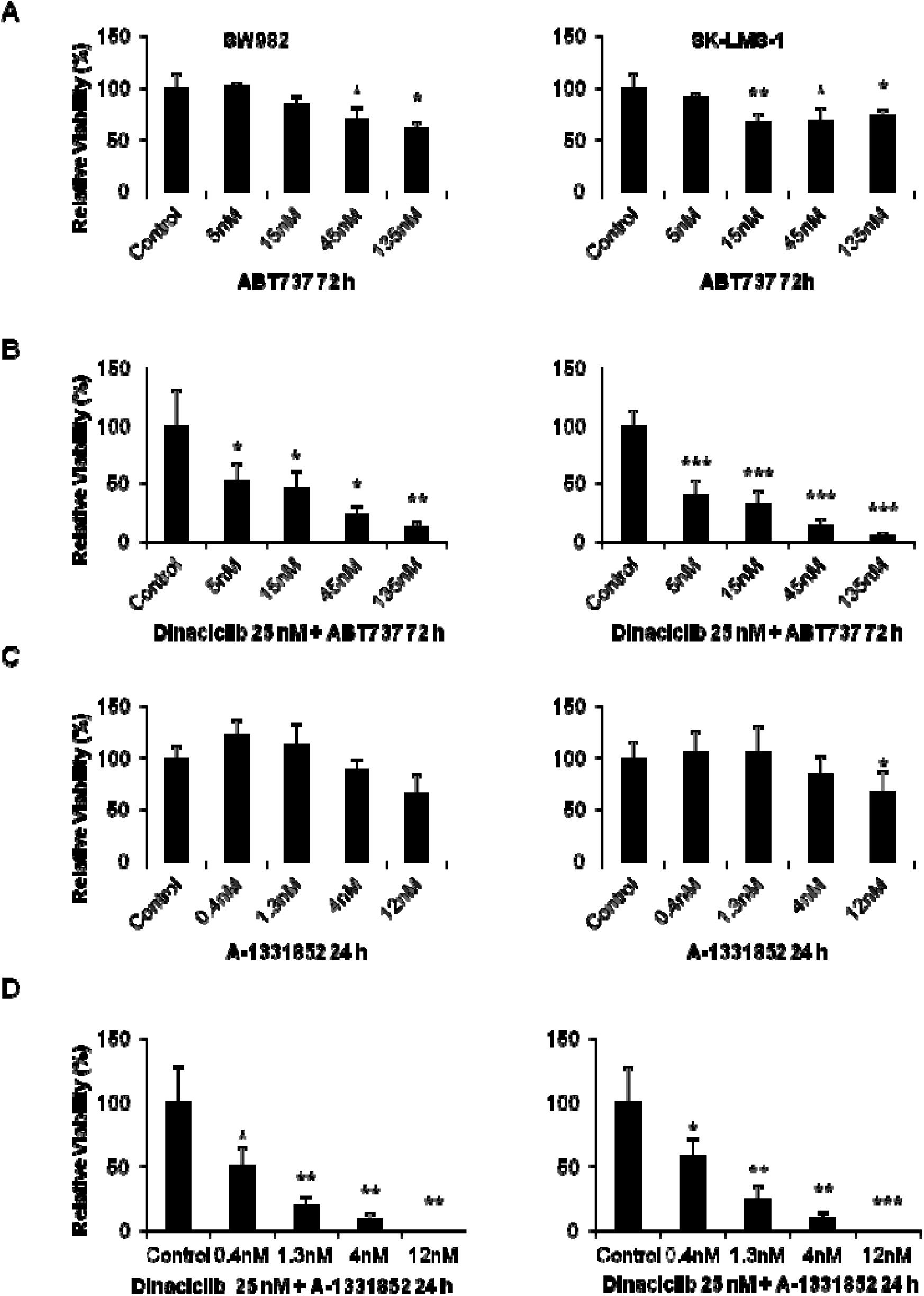
(related to Figure 5)**: BH3-mimetics are unable to induce relevant harm to STS cell lines as monotherapy.** (A to D) Viability measured by means of WST-1 reduction test in SW982 (left) and SK-LMS-1 (right) cell lines after:(A) 72 h treatment with increasing concentrations of ABT-737; (B) 72 h treatment with increasing concentrations of ABT-737 combined with 25 nM Dinaciclib; (C) 24 h treatment with increasing concentrations of A-1331852; (D) 24 h treatment with increasing concentrations of A-1331852 combined with 25 nM Dinaciclib. Data are presented as means ± SD. Statistical significance was achieved by the Student’s *t* test from at least three different experiments: \**p* ≤ 0.05; \*\**p* ≤ 0.001; \*\*\**p* ≤ 0.0001.

**Figure S5:**
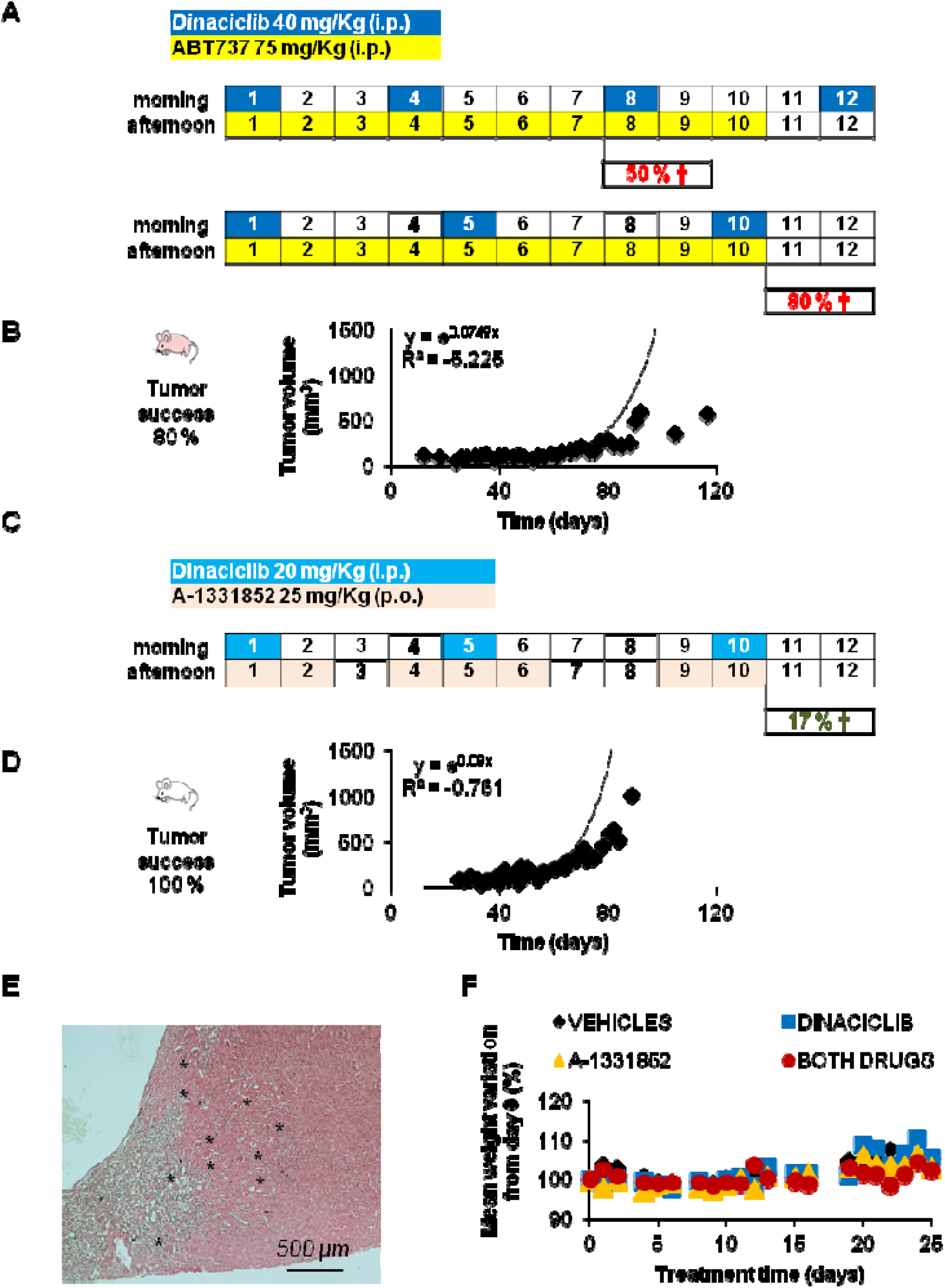
(related to Figure 5)**: Combined treatment with Dinaciclib and BH3-mimetics shows deathly liver toxicity.** (A) Treatment regimes using Dinaciclib and ABT-737 in Hsd:Athymic Nude*Foxn1*^*nu*^ mice. Mortality rates are indicated. (B) Tumor success rate and growth curve for SK-LMS-1 cell line in Hsd:Athymic Nude*Foxn1*^*nu*^ mice. (C) Treatment regime with Dinaciclib and A-1331852 tested in CB17.Cg-*Prkdc*^*scid*^*Lyst*^*bg-J*^/Crl mice. (D) Tumor success rate and growth curve for SK-LMS-1 cell line in CB17.Cg-*Prkdc*^*scid*^*Lyst*^*bg-J*^/Crl mice. (E) Microscopical imaging of H&E staining of one intra-peritoneal sample tumor showing tissue transition from renal cortex to tumor mass. * indicate renal corpuscles. (F) Mean CB17.Cg-*Prkdc*^*scid*^*Lyst*^*bg-J*^/Crl mice weight variation during combination treatment with Dinaciclib and A-1331852. Time 0 represents the beginning of the treatment.

